# Finding coarse and fine scale population structure in a coastal species: population demographics meets genomics

**DOI:** 10.1101/2022.02.15.480495

**Authors:** Diana Catarino, Per Erik Jorde, Lauren Rogers, Jon Albretsen, Marlene Jahnke, Marte Sodeland, Ida Mellerud, Carl Andre, Halvor Knutsen

## Abstract

Population genetic studies often focus on patterns at a regional scale and use spatially aggregated samples to draw inferences about population structure and drivers, potentially masking ecologically relevant population sub-structure and dynamics. In this study we use a multidisciplinary approach combining genomic, demographic, and habitat data with an oceanographic particle drift model, to unravel the patterns of genetic structure at different scales in the black goby (*Gobius niger*) along the Norwegian coast. Using a high-density sampling protocol, we observed restricted gene flow both at a surprisingly fine (kms) and large (100s km) scale. Our results showed a pattern of isolation by distance related to the level of exposure along the Skagerrak coast, where sheltered sampling stations had an overall level of genetic divergence about three times higher (*F*_ST_ =0.0046) than levels observed among exposed samples (*F*_ST_ =0.0015). These results were corroborated by demographic analyses which showed that population-fluctuations decrease in synchrony with distance at much smaller scales for sheltered samples (20 km) than for exposed sites (80 km), suggesting higher population connectivity among exposed sites. We also found a pronounced genetic discontinuity between populations along the Norwegian west and east coasts, with a sharp “break” around the southern tip of Norway, likely driven both by lack of habitat and by oceanographic features.

## Introduction

Many marine species are broadcast spawners and disperse at early life stages by drifting with ocean currents. Traditionally, these species were expected to show little or no genetic population structure across small (kms) or even large geographic scales (ocean regions; i.e. Longmore et al., 2014). Although for some species this holds true, in the past decades this paradigm has considerably shifted (Palumbi, 2004; Hauser & Carvalho 2008), and it has been shown for many species that phenomena such as ocean currents, bathymetry, local adaptation, habitat fragmentation, or reduced migration, limit genetic connectivity across different spatial and temporal levels (i.e. Catarino et al., 2015; Kelley et al., 2016; Barth et al., 2019). In fact, many species with extensive geographic ranges may consist of genetically separated populations despite an apparent homogeneous distribution. Therefore, understanding and identifying the geographic scales of connectivity is a crucial step for basic and applied ecology of marine species (Cowen et al., 2006; Selkoe et al., 2016; Benestan et al. 2021), since this information is essential to improve conservation efforts such as the design of marine reserve networks at suitable scales (Almany et al., 2009; Mertens et al., 2018) and to define stock boundaries for fisheries management (Bernatchez et al., 2017; Quintela et al., 2020; Lindegren et al., 2022). However, detecting the magnitude and geographic scale of connectivity of fish in the marine environment is not always an easy task. Many studies may have missed existing population divergence because population genetic studies usually focus on patterns at a regional scale and use spatially aggregated samples to draw inferences about population structure and drivers, potentially masking ecologically relevant population sub-structure and dynamics. Given that most marine species have wide ranges, spatial sampling for genetic studies is often geographically coarse, since financial and technical constraints limit the number of sampled specimens and their geographical coverage (Martinsohn et al., 2019). Apparent homogeneity over large areas in an initial screening may therefore yield the impression of panmixia and discourage further studies at finer geographic scales. While a coarse-scale sampling strategy may be adequate for uncovering the major population genetic patterns in a species, it risks overlooking finer scale patterns that may hint at population processes of biological relevance for local populations (Barth et al., 2019; Benestan et al., 2021).

Many species, marine or terrestrial, live in habitats that in the context of individual movement and dispersal may be considered two-dimensional, meaning that dispersing individuals may reach neighbouring population(s) along several routes. Theoretical analyses found that in two dimensional habitats, gene flow is particularly effective in counteracting genetic divergence (Kimura & Weiss 1964; Rousset, 1997; Slatkin, 1994), implying that population genetic structure may be weak and difficult to detect in the face of sampling noise (Waples, 1998; Palumbi, 2004). A more powerful approach to detect mechanisms that restrict gene flow is to consider species that occupy a linear, one-dimensional habitat where theory predicts that gene flow is a less potent homogenizing force and where genetic structure should be more pronounced than in a two-dimensional habitat when the total amount of gene flow is equal (Rousset, 1997). In the marine environment species living along the coastline approximate such a linear habitat, in contrast to the open ocean or in more complex seascapes which are more two-dimensional. In linear habitats, limited gene flow caused by habitat discontinuities, environmental drivers, or by geographic distance, is expected to result in more pronounced and thereby more easily detectable genetic structures (Kanno et al., 2011).

Information on genetic structure provides important clues to the pattern and magnitude of gene flow, but genetic data alone are insufficient to determine the demographic effects on populations. While dispersal will leave genetic signatures due to gene flow, the exchange of individuals among locations or local populations can also affect spatial population dynamics, as measured by local or regional variation in abundance through time. Dispersal acts to synchronize population dynamics across space (Bjørnstad et al. 1999), and the spatial scale of population synchrony has been used to delineate population units in marine ecosystems (Östman et al. 2017). Analysis of spatial population synchrony can therefore be used alongside analyses of genetic structure to provide insights to the scales and processes affecting dispersal and population connectivity in a more powerful approach.

In the present study we adopted a multidisciplinary approach to unravel gene flow and population connectivity at coarse and fine geographic scales of a coastal fish species. Using a geographically fine-scaled sampling protocol, we combine genomic data (ddRAD Seq) of the black goby with analysis of spatial population dynamics from abundance time-series, as well as oceanographic modelling of pelagic larval drift, and habitat analyses to identify features that influence local- and regional-scale population connectivity and genetic structure. Specifically, we investigate the effects of scale consideration and sampling design on genetic and demographic inferences and the ecological drivers of the patters observed.

## Material and Methods

### Study Species

The black goby, *Gobius niger* Linnaeus, 1758 is widely distributed in the Eastern Atlantic and Mediterranean Sea, from Mauritania (North Africa) to the eastern Atlantic coast northwards to Norway and into the Baltic Sea (Froese & Pauly, 2021). The black goby is a small coastal fish growing up to 17 cm total length and can be found mostly on sandy and muddy bottoms but also on hard bottom among algae (Moen & Svensen, 2000). The species inhabits bottoms from 2 to 70 m depth but is most common in depths less than 30m. It can inhabit lagoons with brackish water down to 10‰ salinity (Penthon, 2005). Sexual maturity is reached at around 2 years of age, followed by a sedentary benthic adult life (Miller, 1986). Black goby adult males build and protect a nest where the females deposit their eggs. Spawning begins when the water temperature reaches 12°C, and the incubation of the eggs at this temperature lasts for about 20 days (Vaas et al., 1975). The eggs are guarded by the male until they hatch. After hatching, the duration of the planktonic larval phase is approximately 28 days (Planes, 1998). The species has no commercial value, but it is ecologically important in the coastal food web complex as prey for commercially important species such as cod, haddock, and trout (e.g. Hop et al., 1992; Knutsen et al., 2001).

### Study area and sampling

In this study we focused on the Norwegian coastal system from the eastern border with Sweden, along the southern coast of Norway, and up to Ålesund (62° N) in western Norway. Most of the sampling sites are on the Skagerrak coast, which lies between the North Sea and the Kattegat, bordered by Denmark, Sweden, and southern Norway (Figure 1). The Norwegian coast is characterized by an indented landscape with numerous islands, islets and enclosed bays, and cut by several fjords that can extend for several kilometres. The rough coastline and the scattered landmasses create a dynamic and complex oceanography along the coast, but with a general ocean current pattern flowing westwards along the Norwegian Skagerrak coast (the Norwegian Coastal Current, NCC; Albretsen et al., 2012).

**Figure 1.**
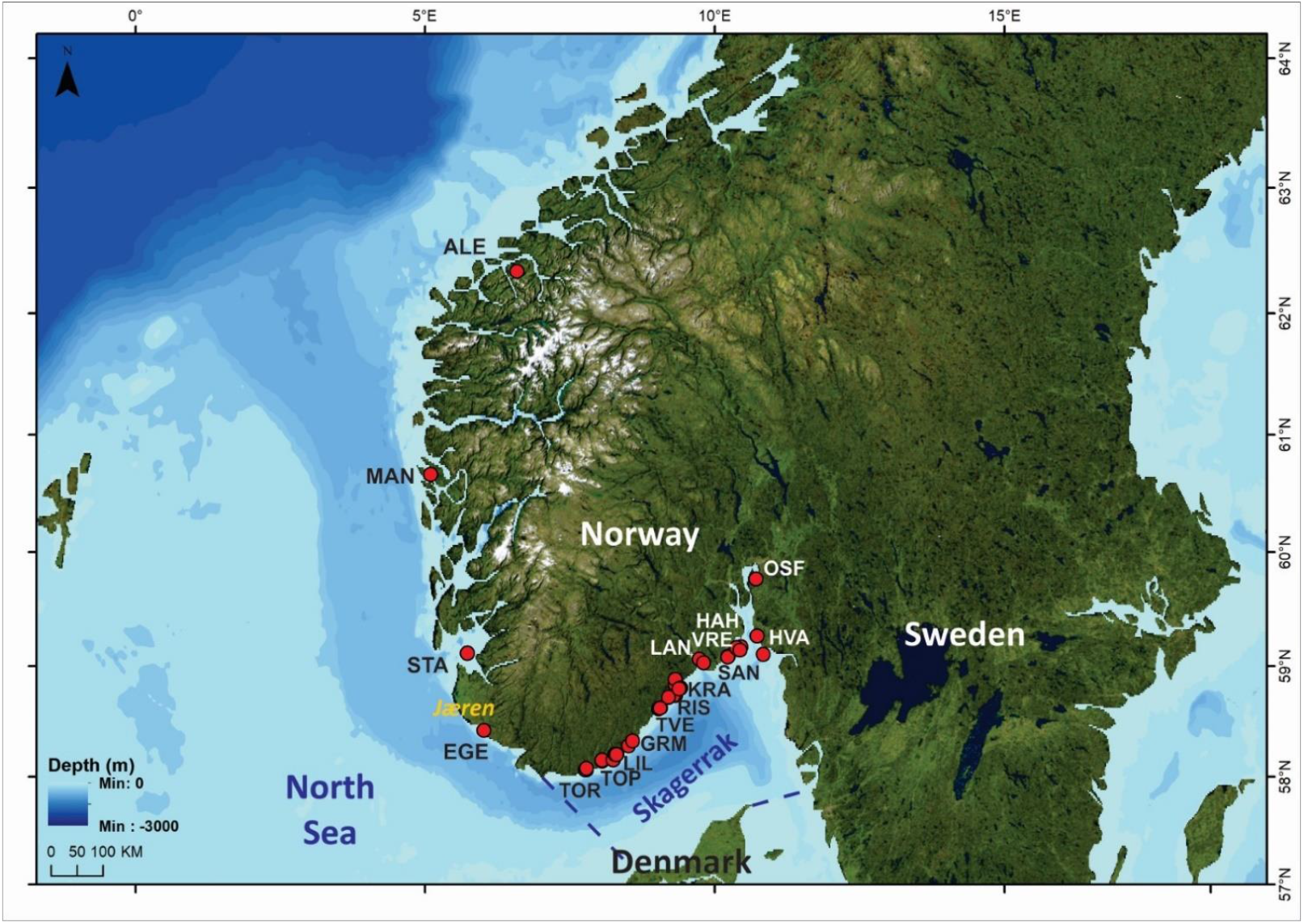
Map showing the main sampling locations where black goby samples were collected for the genetic analyses. *Jæren* denotes the area of a long stretch of sandy beaches. Dashed lines delimitate the Skagerrak region between Norway, Denmark and Sweden. Sampling location abbreviations are given in Table 1.

For more than a century, since 1919, the Institute of Marine Research has conducted an autumn beach seine survey at a set of fixed stations along the Norwegian Skagerrak coast (Stenseth et al., 1999; Table S1). The beach seine is 40 m long, with a stretched mesh size of 1.5 cm, and samples an area up to 700 m^2^ of nearshore (< 15 m depth) habitat (Barcelo et al., 2016; Rogers et al., 2014). In this survey black goby abundance has been counted as number of individuals per haul yearly since 1989 at more than 130 stations distributed among 13 main regions from Torvefjorden (TOR) in the south to the Swedish border in the east (Figure 1; Table1; Table S1). Biogenic habitat type (e.g., eelgrass, green or brown macroalgae) and coverage (scaled from 1-5 subjectively: 1 = bare substrate to 5 = full coverage) have been characterized at each haul station (Table S1). Stations range from sheltered locations inside fjords to exposed areas on the open Skagerrak coast and have been classified as “sheltered” or “exposed” according to their degree of exposure to the open sea (see Rogers et al., 2014).

**Table 1.** Sampling locations and summary statistics for black goby samples along the Norwegian coast (WC, West Coast; SK, Skagerrak). Coordinates are given in decimal degrees (the shown coordinates are an average value when more than one station was sampled). N1 refers to the total number of individuals initially genotyped, combined over sampling stations; N2 is the number of black gobies remaining after filtering and used for downstream analyses (292 in total); Stations referrer to the total number of stations used within a sampling location; expected (He) and observed (Ho) heterozygosity, and inbreeding coefficient (*F*_is_) averaged over 611 SNPs.

| Area | Sampling locations | Site code | Latitude (N) | Longitude (E) | N1 | N2 | Stations | $H_e$ | $H_o$ | $F_{is}$ |
| --- | --- | --- | --- | --- | --- | --- | --- | --- | --- | --- |
| WC | Ålesund | ALE | 62,338 | 6,581 | 23 | 19 | 1 | 0.15<br>5 | 0.157 | 0.017 |
|  | Manger | MAN | 60,664 | 5,095 | 21 | 18 | 2 | 0.15<br>6 | 0.156 | 0.020 |
|  | Stavanger | STA | 59,114 | 5,727 | 24 | 20 | 1 | 0.15<br>2 | 0.149 | 0.047 |
|  | Egersund | EGE | 58,421 | 6,014 | 23 | 18 | 1 | 0.14<br>8 | 0.144 | 0.052 |
| SK | Torvefjorden | TOR | 58,064 | 7,764 | 16 | 14 | 4 | 0.15<br>2 | 0.152 | 0.009 |
|  | Topdalsfjorden | TOP | 58,144 | 8,050 | 24 | 21 | 2 | 0.15<br>5 | 0.155 | 0.024 |
|  | Lillesand | LIL | 58,206 | 8,308 | 24 | 21 | 5 | 0.14<br>6 | 0.148 | 0.004 |
|  | Grimstad | GRM | 58,283 | 8,509 | 17 | 9 | 2 | 0.14<br>5 | 0.150 | -0.036 |
|  | Tvedestrand | TVE | 58,623 | 9,058 | 24 | 17 | 3 | 0.14<br>5 | 0.142 | 0.042 |
|  | Risør | RIS | 58,737 | 9,314 | 24 | 11 | 4 | 0.14<br>1 | 0.150 | -0.038 |
|  | Kragerø | KRA | 58,835 | 9,319 | 24 | 18 | 4 | 0.14<br>3 | 0.139 | 0.046 |
|  | Langesund | LAN | 59,030 | 9,808 | 24 | 17 | 3 | 0.14<br>7 | 0.145 | 0.038 |
|  | Sandefjord | SAN | 59,088 | 10,227 | 13 | 12 | 2 | 0.13<br>9 | 0.137 | 0.023 |
|  | Vrengen | VRE | 59,165 | 10,377 | 25 | 20 | 3 | 0.15<br>3 | 0.146 | 0.056 |
|  | Håholmen | HAH | 59,139 | 10,434 | 24 | 20 | 1 | 0.14<br>4 | 0.138 | 0.054 |
|  | Oslofjord | OSF | 59,766 | 10,712 | 24 | 18 | 1 | 0.14<br>7 | 0.143 | 0.036 |
|  | Hvaler | HVA | 59,264 | 10,731 | 25 | 19 | 3 | 0.14<br>7 | 0.148 | -0.010 |

Samples of *G. niger* for genetic analyses were obtained from a total of 17 different locations (Figure 1, Table 1 and S1) between June and October of 2018. These locations included the 13 main locations described above along the Skagerrak coast (hereafter referred to as SK) and representing 37 hauled stations from the beach seine survey, and 4 additional locations from Egersund (EGE) in the southwest, and along the Norwegian West coast (WC), including in Stavanger (STA), Manger (MAN) and Ålesund (ALE). Fin clips were taken from each fish and stored in 96% ethanol until DNA extraction.

Fish samples were collected in accordance with Norwegian regulations and with all permits required by the Norwegian Government. The studied species (*G. niger*) is not listed in CITES and it is listed as “Least Concern” by the International Union for Conservation of Nature (IUCN; Carpenter et al., 2015).

### Laboratory procedures

Genomic DNA was extracted using the E-Z® 96 Tissue DNA kit (Omega, USA) following manufacturer’s procedures. DNA was quantified on a Fluoroskan FL (Thermo Scientific, USA) using the Quant-iT Broad Range dsDNA assay kit (Thermo Scientific, USA), and integrity was checked by gel electrophoresis on a Multiskan SkyHigh spectrophotometric microplate reader (Thermo Scientific, USA). Double-digest RADseq libraries (ddRAD, Peterson et al., 2012) were prepared using restriction enzymes Sbf1 HF/Sph1 HF. The 95 samples (with 3 replicates) and one blank per plate were digested at 37°C for 45 minutes, and then combined in a ligation mixture with T4 ligase, a general P2 adapter and P1 barcoded adapter and incubated at 16°C for 1 hour 30 minutes, 22°C for 1 hour and 30 minutes and heat inactivated at 65°C for 10 minutes. Successfully ligated samples were size selected (250-600bp) on a Blue Pippin (Sage Science), followed by the AMPure magnetic bead clean-up (Beckman Coulter, USA) and then amplified by PCR: 30s at 98°C, 30s at 98°C followed by 14 cycles of 10 seconds at 98°C, 30 seconds at 65°C, and 30 seconds at 72°C, followed by a final extension of 2 minutes at 72°C. The resulting PCR was cleaned with the AMPure magnetic bead clean-up (Beckman Coulter, USA). Library preparation success was then evaluated by running the libraries on a Bioanalyzer (Agilent, USA), and quantified on Qubit (Thermo Scientific, USA). Libraries were diluted to 14pM, spiked with 5% PhiX and sequenced on an in-house Illumina MiSeq platform (160bp Paired End), by using one lane per plate and the MiSeq v3 reagents kit.

### Bioinformatics and statistical analyses

Sequencing reads for individual specimens were retrieved using the *process_radtags* module in the Stacks software package (Catchen et al., 2013). De-novo assembly of reads from all specimens, as well as variant detection and initial SNP filtering was conducted using the *ustacks, cstacks, sstacks* and *populations* modules in the same software package. An average of 549’980 PE 150bp reads were utilized per specimen in stacks formation. Minimum stack depth (m) for SNP detection was set to 2.

Using VCFtools (Danecek et al., 2011), an initial five different SNPs datasets were generated, each allowing different levels of missing data: 10, 30, 50, 70, and 80%, respectively. Comparing the analyses of these datasets showed that the main genetic pattern found did not change with different levels of missing data. Therefore, the single, final dataset was filtered to retain SNPs with a minimum read depth of 8, a minimum allele count of 2 and a maximum of 50% missing data for locus and individuals. Loci showing deviations of HWE at <0.001 were also removed and the dataset was further filtered to remove loci exhibiting linkage disequilibrium (LD at 0.5 level) with PLINK 1.9 (Purcell et al., 2007). In the end a total of 611 SNPs and 292 individuals were retained for downstream analyses.

Summary statistics, sample sizes, observed and expected heterozygosity, and *F*_IS_ were calculated using the “diveRsity” (Keenan et al., 2013) R package (R Core Team, 2021). Variation in heterozygosity across sites was investigated using a linear model in R. BayeScan (v 2.1., Foll & Gaggiotti 2008) and PCAdapt (Luu, Bazin, & Blum, 2017; Privé et al., 2020) were used to test if loci deviated from neutrality across localities, and between west coast and Skagerrak samples. BayeScan was run by setting sample size to 10,000 and the thinning interval to 50 as suggested by Foll & Gaggiotti (2008). In PCAdapt population structure was first assessed via PCA, and outliers were thereafter detected with respect to how they relate to population structure. The number of principal components (K=2) to identify SNPs deviating from neutrality was then selected by Cattell’s rule (Cattell, 1966). The resulting *P*-values were corrected for false positives using false-discovery rate (FDR) approach at a 0.05 threshold in both cases (Benjamini & Hochberg, 1995). Following a consensus approach, only those loci flagged as outliers by both methods were considered as being under selection (Narum & Hess, 2011).

Pairwise *F*_ST_ (Weir & Cockerham 1984) was calculated using the Genepop R package (v1.1.7, Rousset, 2008), and statistical significance of pairwise *F*_ST_ tests was assessed by G-tests for allele frequency differences, using 10 000 dememorizations and batches, and 1000 iterations per batch. *P*-values in multiple tests were evaluated for significance using the FDR approach. Genetic clustering was investigated by discriminant analyses of principal components (DAPC, Jombart et al., 2010) using the adegenet R package (Jombart & Ahmed, 2011), to provide a graphic description of the genetic divergence among sample locations in multivariate space. The analyses were performed for the whole dataset and for Skagerrak and Egersund separately. The number of principal components kept was determined from the number of PCAs needed to explain approximately 80% of variation. Optimal clustering was retrieved with the function *find*.*clusters* where the best number of clusters is indicated by the lowest Bayesian information criterion (BIC; Schwarz, 1978), inferred by the *k*-means algorithm with 10^6^ simulations.

Isolation by distance (IBD) was analysed for the whole dataset and for Skagerrak only, by testing for a correlation between pairwise genetic *F*_ST_/(1 − *F*_ST_) and geographic distances (overwater distances) using Mantel tests in the Genepop R package, with 10 000 permutations. Overwater distances were calculated using the marmap (Pante & Simon-Bouhet, 2013) and fossil (Vavrek, 2011) R packages, and using bathymetric data (Amante & Eakins, 2009) hosted by NOAA (https://www.ngdc.noaa.gov/mgg/global/relief/ETOPO1/docs/ETOPO1.pdf)

The geographic coarse-scale genetic structure was further characterized by non-linear regression of *F*_ST_ against distance along the coast. The analysis was anchored to the geography by considering only pairwise comparisons with the easternmost location (Hvaler, HVA), and then fitting an exponential (sigmoid) function to those *F*_ST_ and distance measures:

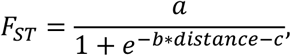

where a, b and c are parameters estimated with the *nls* function in R.

The geographical fine-scale genetic structure of the black goby was characterized using pairwise genetic differences between individuals (Rousset, 2000), as calculated by the Genepop R package (estimator ê). This analysis was restricted to the more densely sampled Skagerrak coast and performed at the beach-seine haul station level. Possible effects of ocean currents on genetic structure were tested by classifying beach seine stations as “exposed” and “sheltered” with respect to the coastal current, following Rogers et al. (2014), and carrying out the analysis separately for pairs of individuals from sheltered and exposed locations (Table S1, Figure S1). The geographic distance between individuals were calculated as distance overwater between beach seine stations. Genetic differences (ê-estimates) for pairs of individuals from sheltered stations were then regressed (linear regression, *lm* function in R) against geographic distance, and similar for exposed pairs. The slopes of the regressions were tested for significance with permutations tests. The tests randomly permuted the genetic distance (ê) matrices repeatedly (10000 times) and regression slopes were calculated for the permuted data. *P*-values were calculated as the fraction of permutations that resulted in a slope that was as large or larger than the observed one.

### Spatial population synchrony and decorrelation scale

Spatial population synchrony, defined as the spatial covariation in population density fluctuations (Bjørnstad et al., 1999), was calculated by taking the pairwise correlation in natural log-transformed catches among each pair of sampling stations, and relating pairwise correlations to the distance between stations. A constant of 1 was added to each count to avoid taking the logarithm of zero. Comparisons were limited to those stations sampled a minimum of 15 years and having at least 10 years with positive (non-zero) catches (n = 132 stations).

An exponential decay model (Bjørnstad et al., 1999) was fitted to pairwise correlations by distance, measured as distance over water between stations:

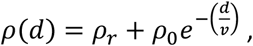

where *ρ*(d) is the pairwise correlation at distance d, ρ_r_ is the asymptotic correlation, or background regional correlation in recruitment, and ρ_r_ + ρ_0_ is the estimated correlation at zero distance, following Rogers et al. (2014). The parameter v (“decorrelation scale”) describes distance at which the pairwise correlation between time series is reduced to ∼37% (e^−1^) of that at zero distance, relative to the background regional level of correlation. To test the hypothesis that abundance at exposed stations would covary over longer distances than sheltered stations due to increased connectivity, models were also fit separately to pairs of exposed stations and pairs of sheltered stations. Confidence intervals (90%) were estimated based on 10000 bootstrap replicates, following Bjørnstad et al. (1999).

### Habitat associations

Habitat associations for black goby were investigated using the Skagerrak beach seine time-series (Table S1). Specifically, we tested for effects of habitat type and percent cover on abundance of black goby in beach seine hauls using a negative binomial generalized mixed effects model which included a random intercept by station. Data from all stations surveyed for at least 15 years (n = 137) were used in this analysis, for a total of 3930 individual observations of black goby abundance. Although recorded as a categorical variable with five categories (“bare substrate” to “full cover”), coverage was treated in the model as a continuous variable (ranging from 1 to 5) after preliminary model runs indicated an approximately linear relationship between abundance and coverage. Models were fitted using the lme4 package (Bates et al., 2015) in R.

### Oceanographic modelling

A combination of high-resolution hydrodynamical modelling and particle-tracking was applied to investigate dispersal potential of larvae from all beach seine locations, and additional sites around the southern tip of Norway, and in the west encompassing the genetic sampling stations between Egersund and Stavanger (see Table S1). Our circulation model system consisted of three high-resolution fjord models run in parallel with input along the open boundaries from the Norwegian coastal model NorKyst800 (Asplin et al., 2020). The highest resolution models all applied 160m resolution in the horizontal, and similar experiments and validation analysis are presented in e.g. Dalsøren et al., 2020. We applied the open-source Regional Ocean Modeling System (ROMS) (Haidvogel et al., 2008; Shchepetkin & McWilliams, 2005) (see also http://myroms.org), which is a state-of-the-art, three-dimensional, free-surface, hydrostatic, primitive equation ocean model that uses generalized terrain-following s-coordinates in the vertical. The results from the three model grids using 160m resolution covering the inner Skagerrak, the southernmost part of the Norwegian coast and the southwestern fjords, respectively, were merged into one large grid before being used as input in the particle-tracking model.

Our drift simulations of larvae-particles were conducted by using the open access Lagrangian Advection and Diffusion Model (LADIM; https://github.com/bjornaa/ladim). The model solves the Lagrangian equation of motion using a 2nd-order Runge-Kutta scheme. Due to limited knowledge on larval swimming behaviour, we initialized the particles in 1, 2, 3 and 4m depth and left them without any vertical movement. 250 particles were released in each depth at all 204 locations which gives 1000 particles per station. To include slightly different physical environments, we performed three identical simulations initiating June 2^nd^, July 1^st^ and August 1^st^, all in 2018 as a representative year, and followed every particle for 25 days. A small horizontal random walk was also implemented to ensure variability between the particles and to represent small-scale turbulence.

## Results

### Population genetic patters

The final dataset consisted of 292 individuals from 17 main localities genotyped for a total of 611 SNPs, with no deviations from HWE or LD. Genetic variability (expected heterozygosity, H_e_) ranged from 0.139 for SAN locality to 0.156 at MAN (Table 1) and displayed a significant increasing trend from east to west (linear regression of H_e_ on distance over water from the easternmost locality, Hvaler: slope = 1.285*10^−5^-; R^2^ = 0.3731; *P* = 0.0055).

BayeScan identified three possible outlier loci out of the 611 SNPs analysed for the whole dataset, while no outlier loci were identified when comparing West Coast *vs*. Skagerrak. PCAdapt identified a total of 15 outlier loci. Following a consensus approach, two SNPs (SNP 9075_98 and 13326_52) were identified by both methods as outlier candidates under selection. Pairwise *F*_ST_, DAPC and IBD analysis were performed both with and without the two candidate outlier loci and since no significant differences were observed in the results (Figure S2), all loci were retained for all analyses presented below.

Cluster analyses (DAPC) comprising the whole dataset clearly showed two main clusters separated along DF1 (Figure 2a), one grouping together the West Coast samples (ALE, MAN and STA), and a second grouping all Skagerrak samples and Egersund. This result is supported by the lowest BIC, which indicated that *K*=2 was the most likely number of clusters (Figure S3). Using only Skagerrak and Egersund samples in a separated DAPC analyses (Figure 2b), no further pattern stood out with samples showing no clear separation, and many locations overlapping. This genetic pattern is also supported by *F*_ST_ analyses, where significant genetic differentiation was found for overall sampling localities (*F*_ST_ overall =0.0181, *P*<0.001). Pairwise *F*_ST_ values ranged from -0.0065 to 0.0657 averaged over the 611 SNP considered, with the highest values observed for the pairwise comparisons between West Coast localities and Skagerrak/Egersund, which were highly significant (Table 2; overall WC *vs*. SK/EGE *F*_ST_= 0.0400, *P*<0.001). In contrast, pairwise comparisons showed no significant differences among West Coast localities and Stavanger nor among Skagerrak and Egersund (Table 2). Analysis of IBD showed that for the entire Norwegian coast, genetic divergence *F*_ST_/(1−*F*_ST_) increased with increasing distance (Figure 3a; Mantel, *P* <0.001), while no such relationship was observed within Skagerrak samples only (Figure 3a, Mantel, *P*>0.05). This suggests that the divergence pattern found is mostly driven by the West Coast samples and is illustrated with the sigmoid curve in Figure 3b, evidencing a sharp increase in *F*_ST_ around the southern tip of Norway, located between the EGE and STA localities.

**Table 2.**
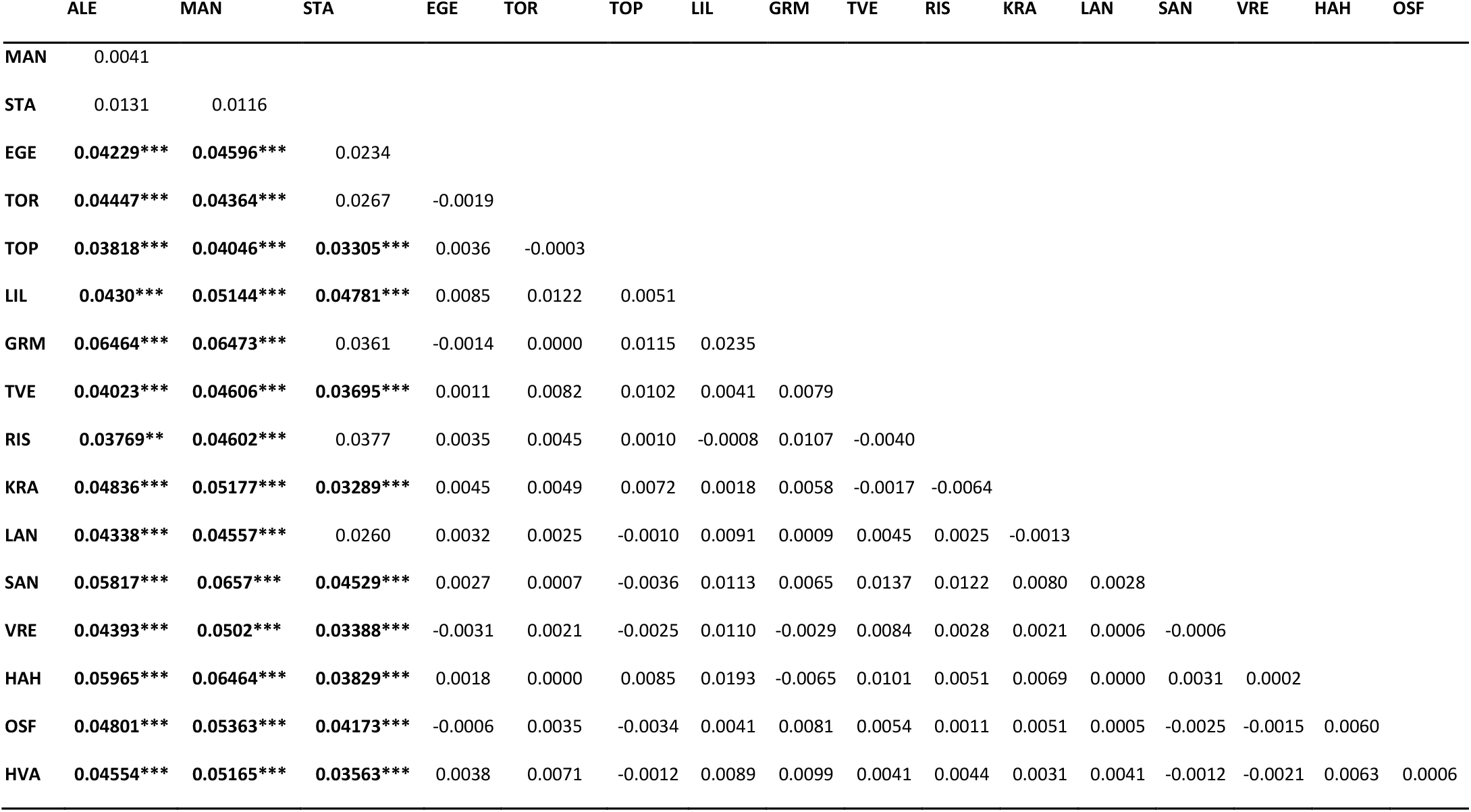
Pairwise *F*_ST_ among sampling locations with significant estimates in bold and level of significance after FDR correction denoted by asterisk: * p<0.05; ** p<0.01; *** p<0.001. See Table 1 for sample abbreviations.

**Figure 2.**
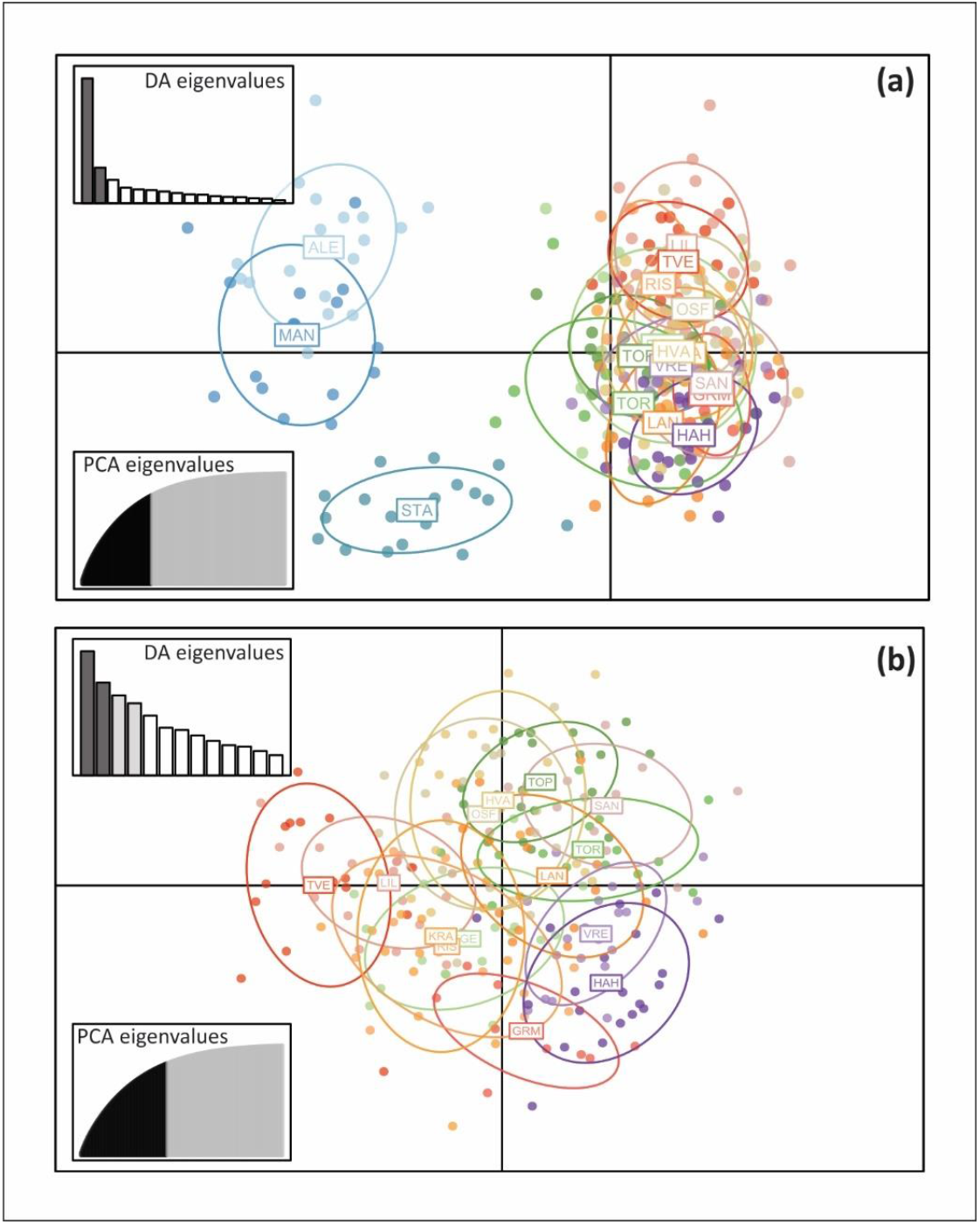
Results from the discriminant analysis of principal components (DAPC) showing (a) two clusters, one corresponding to West coast locations and the other to Skagerrak+ Egersund samples; and showing (b) no distinct clustering within Skagerrak +Egersund only. The DAPC was conducted based on 100 PCs that conserved more than 80% variation.

**Figure 3.**
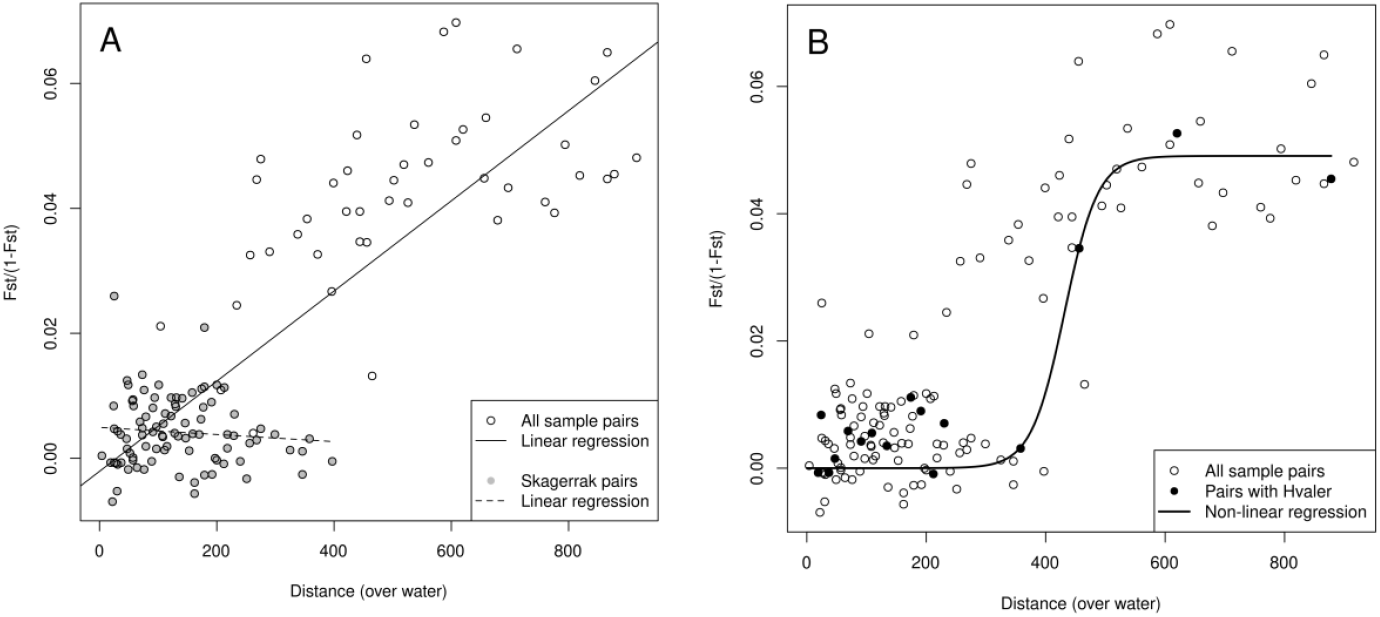
Isolation by distance (IBD) for the black goby. Effect of geographic distance (km over water) on linearized observed genetic divergence, *F*_ST_/(1-*F*_ST_), (a) among all black goby sampling localities (slope = 7.2*10^−5^, *P*<0.001) and for Skagerrak localities only (slope= 3.8*10^−6^, *P*>0.05); (b) among all pairwise comparisons with Hvaler in bold (eastern most sample) and by fitting a sigmoid curve to the data points including Hvaler highlighting the sharp increase in *F*_ST_ from the east to the west coast of Norway.

### Fine-scale genetic structure patterns

Although pairwise comparisons among Skagerrak sampling localities were non-significant (Table 2), and low levels of genetic divergence were found among the Skagerrak sampling localities across all loci, the overall divergence was nevertheless highly significant (overall *F*_ST_=0.0040, *P*<0.001), suggesting the possibility of some cryptic genetic structure within Skagerrak. To look for fine-scale patterns of genetic divergence within Skagerrak, this dataset was analysed separately for IBD among individuals. Although the Skagerrak samples did not show any patterns of IBD when using the whole dataset with the individuals grouped in main sampled localities (Figure 3a, *P*> 0.05; Table 1), when separating according to the level of exposure for each station, two different patterns of genetic differentiation within Skagerrak emerged. First, the overall level of genetic divergence among sheltered samples was about three times higher (overall sheltered *F*_ST_ =0.0046) than among exposed samples (overall exposed *F*_ST_ =0.0015). Second, genetic divergence increased significantly with distance for sheltered stations (Figure 4; *P* = 0.0018; slope = 3.446*10^−5^), but not for exposed stations (Figure 4; *P* = 0.3555; slope = 2.298*10^−6^).

**Figure 4.**
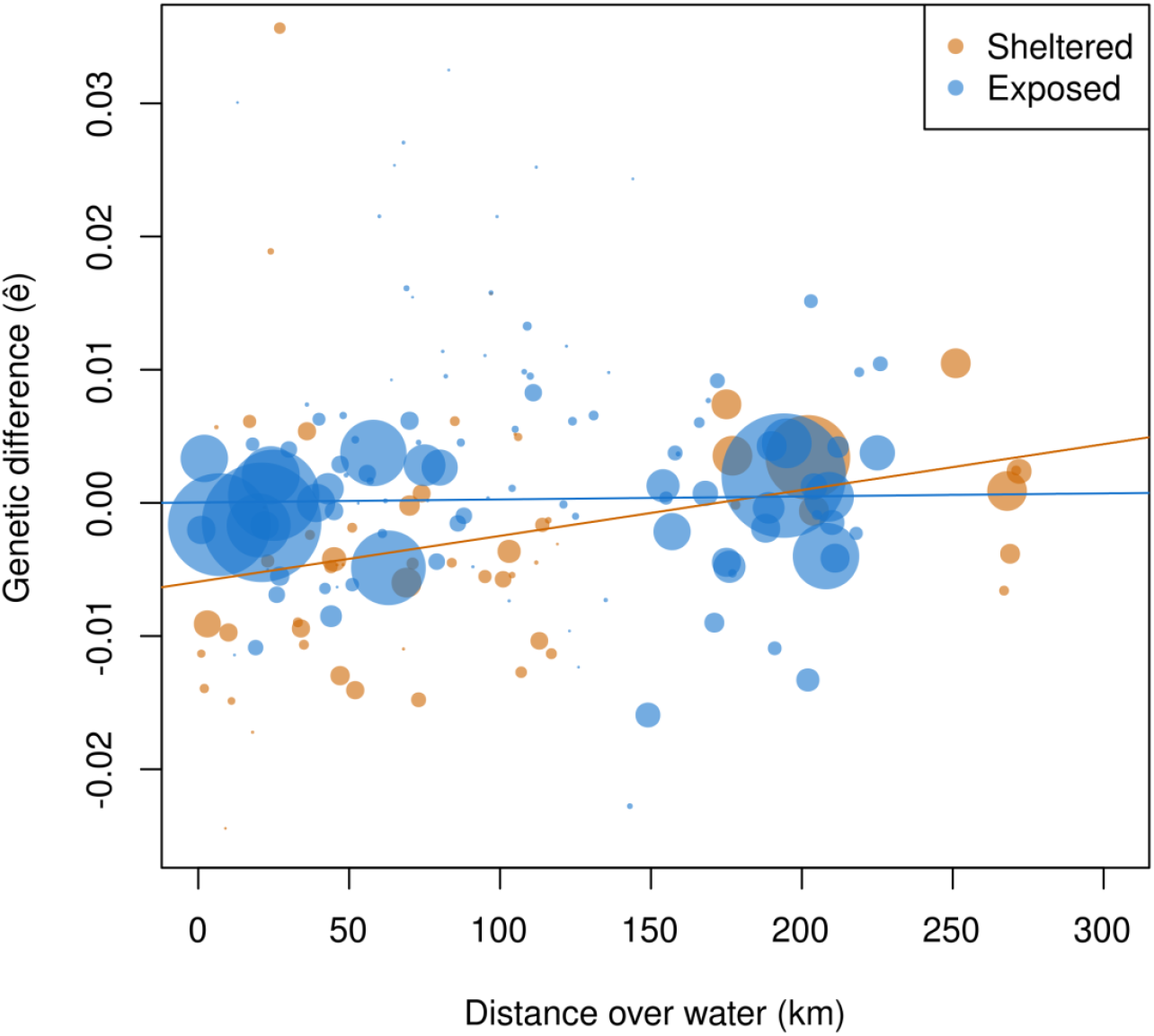
Isolation by distance (IBD) among individuals for the black goby sheltered (orange) and exposed (blue) sampling stations in Skagerrak. Sampling stations were binned into distance classes of 1 km and size of the dots were scaled to number of individual pairs within each distance class. Effect of linear distance on the genetic difference (ê) calculated at individuals’ level, showing a significant increase with distance for sheltered stations (P = 0.0018, slope = 3.446*10^−5^) and non-significate trend for exposed stations (P = 0.3555, slope = 2.298*10^−6^).

### Spatial decorrelation scale

Station-specific fluctuations of black goby abundance in the Skagerrak were positively correlated (i.e., synchronous) at nearby stations, with synchrony decreasing with increasing distance between stations (Figure 5). The spatial decorrelation scale (v) considering both exposed and sheltered stations was estimated as 27.9 km (90% CI = 13.8 -94.8 km). When only sheltered stations were considered, the spatial decorrelation scale was 20 km (90% CI = 2.9 – 266 km), which was substantially smaller than the decorrelation scale among exposed stations of 81 km (90% CI = 15.1 – 254 km), although confidence intervals were overlapping.

**Figure 5.**
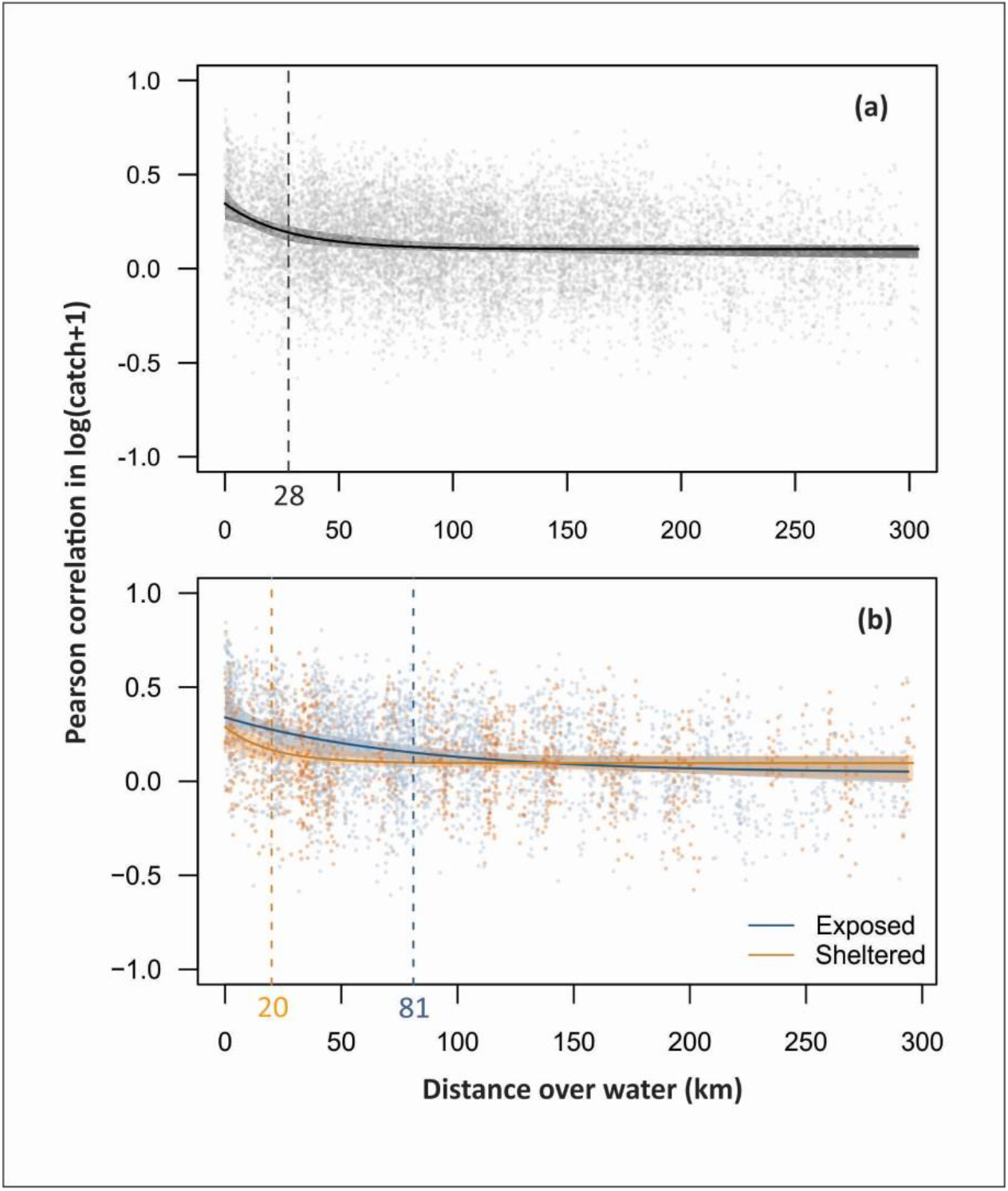
Spatial decorrelation in abundance fluctuations of black goby at beach seine stations along the Skagerrak coast. Results are shown for (a) all stations and (b) models fit to sheltered (orange) and exposed (blue) stations separately. Points show pairwise correlations among stations, and solid lines show a fitted exponential decay model along with bootstrapped 90% confidence intervals (shaded areas). The estimated decorrelation scale (v) is shown as a vertical dashed line.

### Habitat associations

The best model for black goby abundance in the Skagerrak beach seine survey included effects of biogenic habitat type and cover. Black goby catches significantly varied by habitat type (X^2^ = 44.2, df = 5, *P* < 0.001), with highest catches in eelgrass and mixed eelgrass/brown macroalgae habitats, and lowest catches in habitats lacking eelgrass, but containing brown macroalgae or green algae (Figure 6). Black goby catches were also significantly associated with habitat cover, with higher catches at sites with higher coverage percentage (z = 8.42, *P* < 0.001).

**Figure 6.**
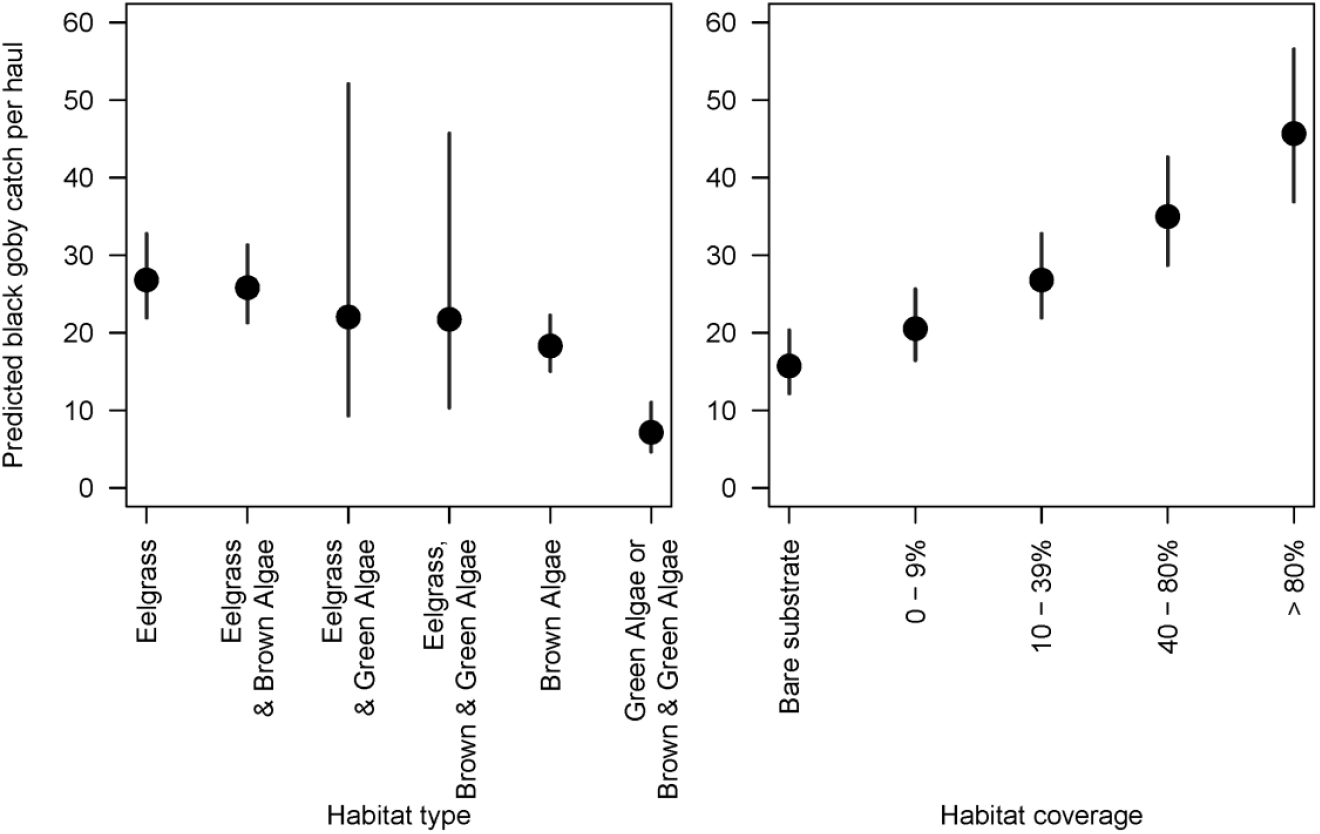
Predicted effects of biogenic habitat type and percent coverage on catches of black goby in the Skagerrak beach seine survey (with 95% prediction intervals). Predictions are shown for intermediate coverage (10-39%) in the first panel, and for eelgrass habitat in the second panel.

### Oceanographic modelling

The oceanographic modelling shows that the larvae-particles released in the surface layer inshore and along the Skagerrak coast, from the Swedish border in the East to Egersund in the West, are subject to three different destinies (Figure 7). (1) A portion is mixed and spread widely by the NCC, which has a persistent westward flow out of the Skagerrak along the Norwegian coast. The NCC in the Skagerrak is relatively strong and often has a distinct core close to the coast. (2) Most larvae-particles entering the NCC have a stronger offshore drift and were not able to settle along the coast at all. (3) A significant and important portion of the larvae are retained locally and hatched inside the fjords, suggesting local settlement. The probability for onshore settlement West of Egersund from larvae-particles released East of Egersund is very small, since the NCC around the southern tip of Norway, transports the drift particles offshore, towards deeper water. For larvae-particles released East of the southern tip of Norway, the NCC acts slightly different with a broader core and occasional long-periodic eddies interfering the continuous transport northward. The localities around Stavanger provide the area with some larvae retention even though most of the larvae-drift from the localities West of Egersund is offshore.

**Figure 7.**
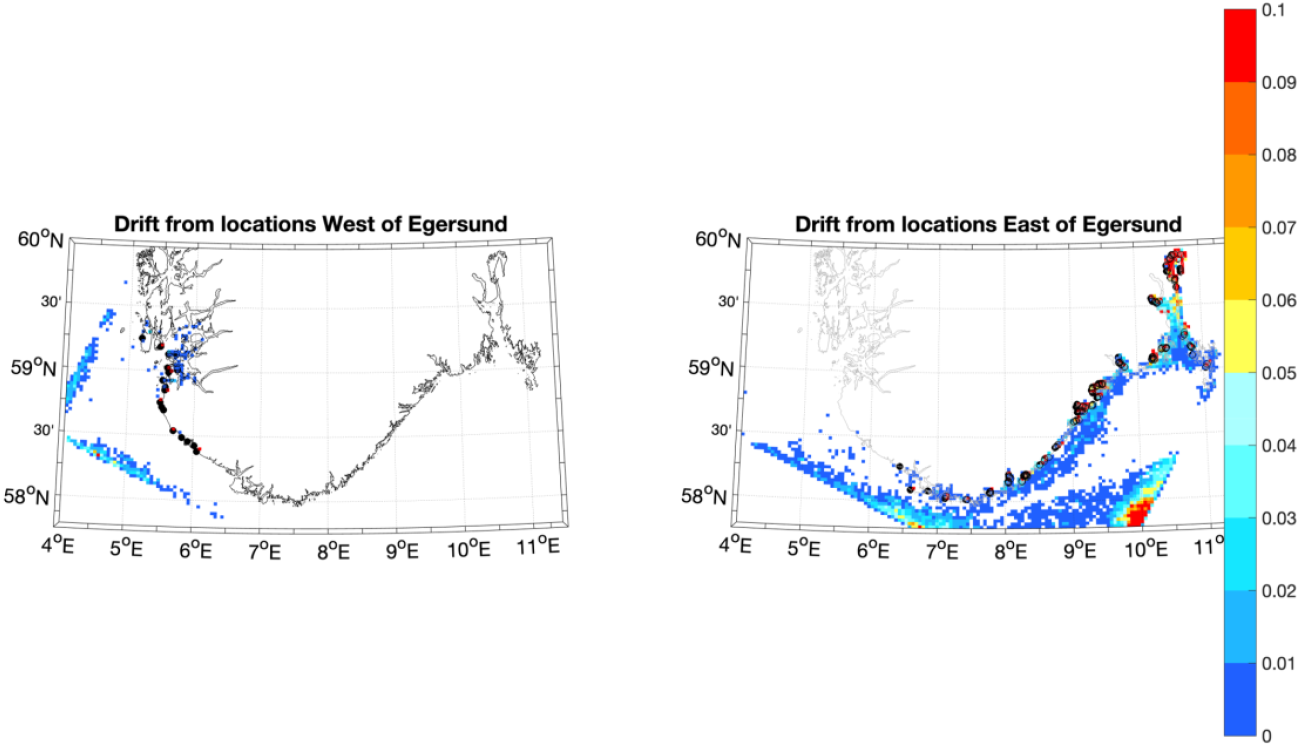
Maps from the oceanographic drift model showing the density of particles (colours) after 25 days drift from the locations West (left panel) and East of Egersund (right panel). The locations where particles were released are denoted with black circles. The colour scale denotes probability in % relative to the total number of particles released from all locations East and West of Egersund. The broad, zones of particle densities offshore are linked to the open boundary of the high-resolution fjord models that inhibits further drift offshore.

## Discussion

Using a multidisciplinary approach, we combined genetic, demographic and habitat data from a long-term data series with an oceanographic particle drift model to unravel the patterns of genetic structure on different geographic scales in the black goby along the Norwegian coast. While geneflow and weak population structure is expected for marine species with high dispersal ability, we observed restricted gene flow both at a surprisingly fine (kilometres) and large/coarse (100kms) scale. Our results show a pattern of isolation by distance along the Skagerrak coast and a pronounced genetic discontinuity between populations along the Norwegian south and west coasts, likely driven by both lack of habitat and oceanography.

Our sampling design focused on the Skagerrak coast, where time series for annual abundance are available, giving the opportunity to investigate fine-scale spatial population patterns both at the genetic and demographic level. At a first glimpse, the general overall pattern showed high levels of connectivity across Skagerrak for both genetic and demographic data. In the Skagerrak, we found no genetic differentiation for pairwise comparisons among locations (Table 2), no IBD, and no clusters were perceived in the DAPC analyses. This is in accordance with the predictions of the oceanographic model where the Norwegian coastal current boosts passive dispersion of larvae promoting connectivity. However, although the overall level of genetic divergence among Skagerrak sample localities was low, it was statistically significant (SK_overall_ *F*_ST_ =0.0040, *P*<0.001), suggesting possible hidden population genetic structure. The dense, fine-scale sampling design allowed us enough spatial resolution to examine samples at the individual level and compare stations that are largely sheltered from the coastal current (i.e., located inside small fjords or bays, or behind skerries) and more exposed areas. On the supposition that ocean currents transport pelagic larvae along the coast, we expect that exchange of larvae will be higher among the more exposed localities and, further, if this current-driven larval dispersal is the dominant mechanisms for gene flow in the species, that gene flow should also be higher among exposed localities than among more sheltered ones. These expectations were indeed met, as the genetic IBD analyses yielded a steeper slope for sheltered than for exposed localities (cf. Figure 4), indicating less gene flow among sheltered localities.

The observed spatial correlation in abundance, or population synchrony, corroborates the genetic findings, as has been shown for other coastal species (Östman et al., 2017). First, nearby stations only a few km apart displayed higher correlations in abundance than stations situated farther apart on the coast (cf. Figure 5). This finding gives evidence that nearby populations are indeed connected demographically, as expected if they also exchange genes. Second, the sheltered stations displayed a more rapid decline in correlation with distance, i.e. a shorter “decorrelation” scale, than did the exposed areas (20 and 81 km, respectively), indicating that dispersal is more restricted among sheltered areas. This finding is also in agreement with genetic data, suggesting a higher level of connectivity among exposed stations than sheltered ones. Our particle drift model showed some level of particle larvae retention, mostly among some of the Skagerrak most enclosed fjords (see Figure 7) supporting lower levels of connectivity or dispersal with outside areas. Nevertheless, power at such small spatial scales was low to test the findings at the demographic and genetic level, and more information of larvae dynamics would be necessary before attempting to model very fine scale dispersal on a nest scale. Still, in the light of the results obtained, it is plausible that exposed stations are more affected by the Norwegian coastal current promoting higher rates of dispersal translating in a higher level of population homogenization, while sheltered areas likely are less influenced by the NCC and experience localized retention processes (i.e. Ciannelli et al., 2010; Myksvoll et al., 2014; Hansen et al., 2021). This correspondence between genetic and demographic patterns is relevant for the long-standing discussion about the use of genetic data for inferring connectivity among natural populations (Lowe & Allendorf 2010; Knutsen et al., 2011), and yields support to the notion that genetic markers provide information of ecological as well as evolutionary relevance.

At a larger scale, when considering the entire study area, we found a strong genetic discontinuity around the southern tip of Norway, between the two sample localities Egersund and Stavanger (cf. Figure 1). We identified two distinct genetic clusters, one comprising the west coast samples and the other comprising all samples from Egersund and the Skagerrak coast, with increased levels of pairwise divergence (*F*_ST_ =0.0400) between West coast and Skagerrak samples. Lower levels of genetic diversity were found within the Skagerrak samples, suggesting restricted geneflow between these two regional groups. Genetic discontinuities, or “genetic breaks,” are characterized by rapid increase in genetic divergence over short geographic distances (Sotka et al., 2004; Catarino et al., 2015). This particular genetic break, around the southern tip of Norway, has been previously identified for other fish species, such as corkwing wrasse (*Symphodus melops*; Knutsen et al., 2013; Blanco Gonzalez et al., 2016; Mattingsdal et al., 2020) ballan wrasse (*Labrus bergylta*; Seljestad et al., 2020), and even for some kelp species (Evankow et al., 2019). The area around this break point, while not exactly localized for any species, is characterized by a bare stretch of sandy beaches (Jæren, about 26 km; Blanco Gonzalez et al., 2016), between Egersund and Stavanger, probably resulting in a habitat discontinuity for some species. For wrasses, hard bottom substrate is an essential habitat, required to build nests where females lay their eggs (Darwall et al., 1992; Villegas-Ríos et al., 2013). Furthermore, telemetry studies showed that wrasses have small home ranges and individual movements are likely to be restricted to less that 300m (Halvorsen et al., 2021), suggesting that individual migration across Jæren is unlikely. For the black goby, not much is known about individual movements, although they share similar life history traits with wrasses: adult black gobies are mostly sedentary, and males are also territorial, building and guarding nests where females lay their sticky eggs. Therefore, for both, black goby and wrasses, dispersal is assumed to be mostly promoted by the pelagic larvae (Knutsen et al., 2013; Kara & Quignard, 2019). Regarding habitat, the black goby has been described in the literature as preferring soft bottoms and unvegetated areas (Perry et al., 2018), and with strong habitat selection shifts based on the presence of predators (Kruschel & Schultz, 2011). However, although black gobies are occasionally captured in bare substrates, our habitat analysis clearly showed that the black goby is more abundant in areas with higher vegetation coverage and prefers eelgrass and mixed eelgrass/brown macroalgae habitats (see Figure 6). This suggests that habitat discontinuity may also be a good explanation for the lack of gene flow across the bare stretches of sandy beaches at Jæren for the black goby population, similarly to what has been found for wrasse species (Blanco Gonzalez et al., 2016; Seljestad et al., 2020). Habitat discontinuities have been also shown to be a major population subdivision in different environments and across different taxa (Riginos and Nachman, 2001; Johansson et al., 2008; Binks et al., 2019) and therefore could be a plausible explanation. Nevertheless, habitat discontinuity alone does not explain how a species with a long pelagic larval stage (about 28 days) is unable to drift over a few dozens of kilometres of the stretch of sandy beaches. Our oceanographic model suggests that particles have a high dispersal potential and therefore larval transport should be high enough to overcome the physical barrier of lack of habitat across Jæren. The results obtained with the oceanographic drift model offers a plausible picture of the drivers promoting the genetic break and restricted geneflow, by showing that particles that are released near the southern tip of Norway drift offshore towards deep water, where recruitment of juveniles seems unlikely, restricting the particles exchange between east and west of the southern. This pattern is driven by the Norwegian coastal current, which flow westwards along the Skagerrak coast (Albretsen et al., 2012), but drifts offshore at the southern tip of Norway. The oceanographic particle drift model also shows that particles released close to Stavanger and Jæren will continue westwards and will not mix with Skagerrak, preventing genetic connectivity and likely strengthening the barrier effect. The model shows that few particles/larvae released from the western end of Skagerrak will reach the Egersund area and this is likely sufficient to homogenise (probably together with some individual migration) the local gene pool, making Egersund more genetically similar to Skagerrak samples. The structuring effect of this oceanographic feature in species with early life stages is also corroborated when comparing with species without such stages. A recent study showed that for the broadnosed pipefish (*Syngnathus typhle*), which gives birth to live young, this barrier effect is less pronounced, and that genetic divergence is likely due to habitat effect and isolation by distance (Knutsen et al., 2022).

Our multidisciplinary approach and fine-scale sampling design allows for detecting the importance of retention of early life history stages in shaping the genetic structure at finer geographic scale in coastal waters. Further, oceanographic circulation also seems to play an important role in the creation of the genetic divergence observed between West coast and Skagerrak samples. Therefore, our results using these combined approaches, indicated a more complex scenario of gene-flow restriction suggesting a combination of both habitat discontinuity but also an important oceanographic component.

## Acknowledgments

This study was funded by the ECOGENOME project (grant nr. grant number: 280453), through the Norwegian Research Council. Further funding was provided by MarGen II, an Intereg project under the Øresund-Kattegat-Skagerrak program, by the Ministry of Trade, Industry and Fisheries, and from University of Agder, Centre for Coastal Research. We thank the scientific team participating in the Beach seine survey. We also thank Ricardo Medeiros (IMAR, University of the Azores) for making the map and to Karen Martinez-Swatson (IMR, Norway) for helping with the laboratory methodology description. DC was funded by the University of Agder and at later stage by FCT – Foundation for Science and Technology, I.P., under the project UIDP/05634/2020 granted to Okeanos. Okeanos is funded through the FCT, I.P., under the project UIDB/05634/2020 and UIDP/05634/2020 and by the Regional Government of the Azores through the initiative to support the Research Centers of the University of the Azores and through the project M1.1.A/REEQ.CIENTÍFICO UI&D/2021/010.

