## Supplementary material for "Finding coarse and fine scale population structure in a coastal species: population demographics meets genomics": Figure S1; Figure S2; Figure S3

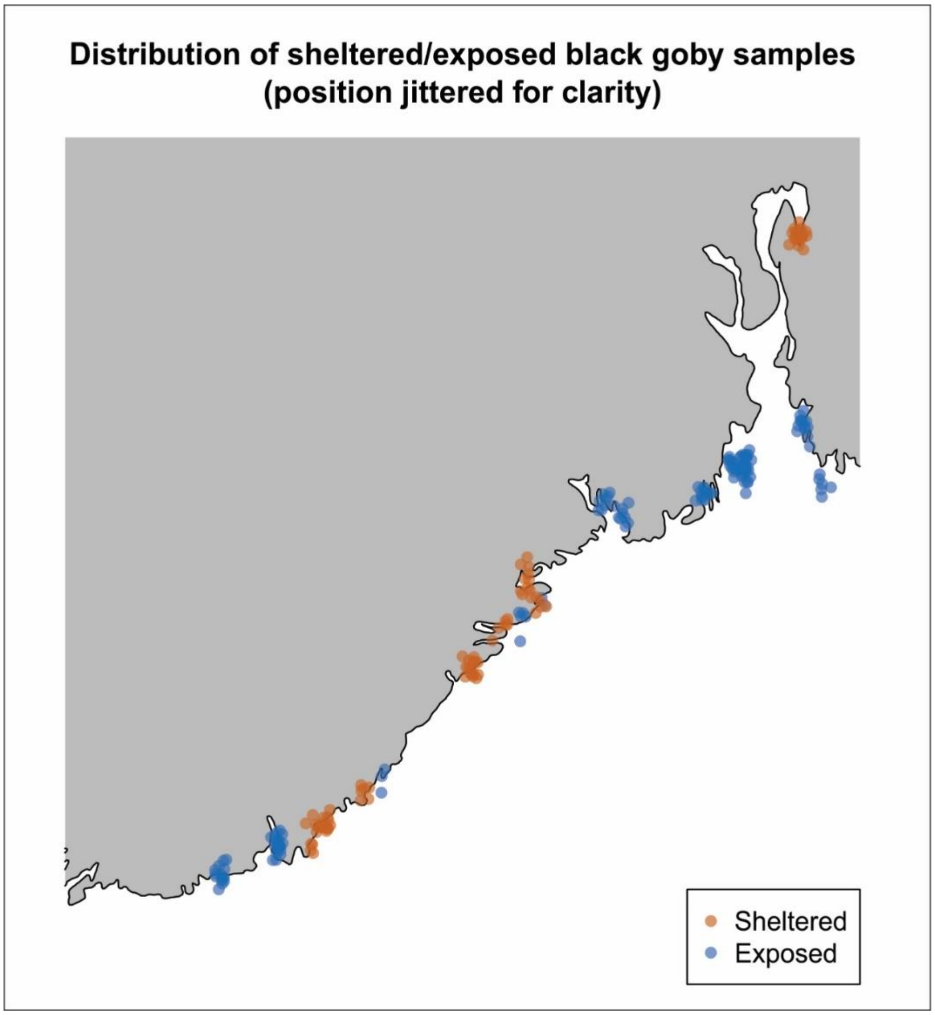


**Figure S1**. Map showing the distribution of the beach seine data stations classified either as sheltered (orange) or exposed (blue) used in isolation by distance and decorrelation analyses.

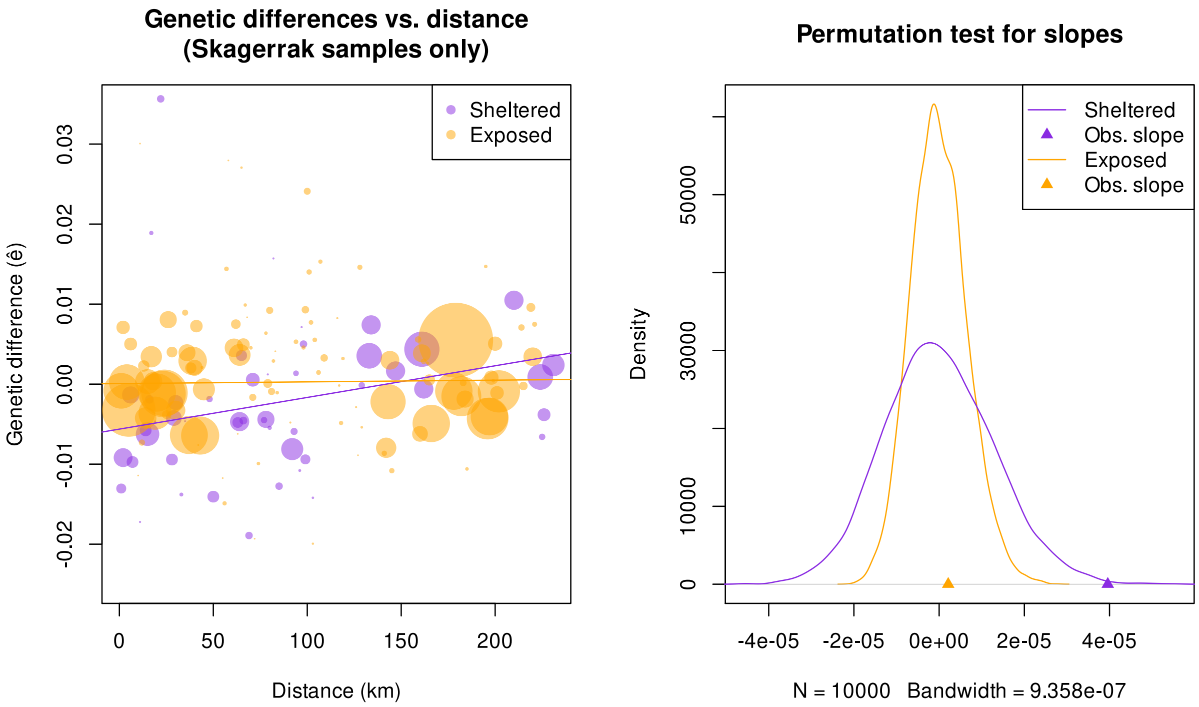


**Figure S2.** Isolation by distance (IBD) and permutation test for the black goby sheltered (purple) and exposed (orange) sampling stations in Skagerrak, without the two outlier loci (SNPs 9075_98 and 13326_52). Effect of linear distance on the genetic difference (ê) calculated at individuals' level. For the sheltered locations P=0.0025, and for the exposed P=0.35, showing no changes in the general patter when the two loci are removed.

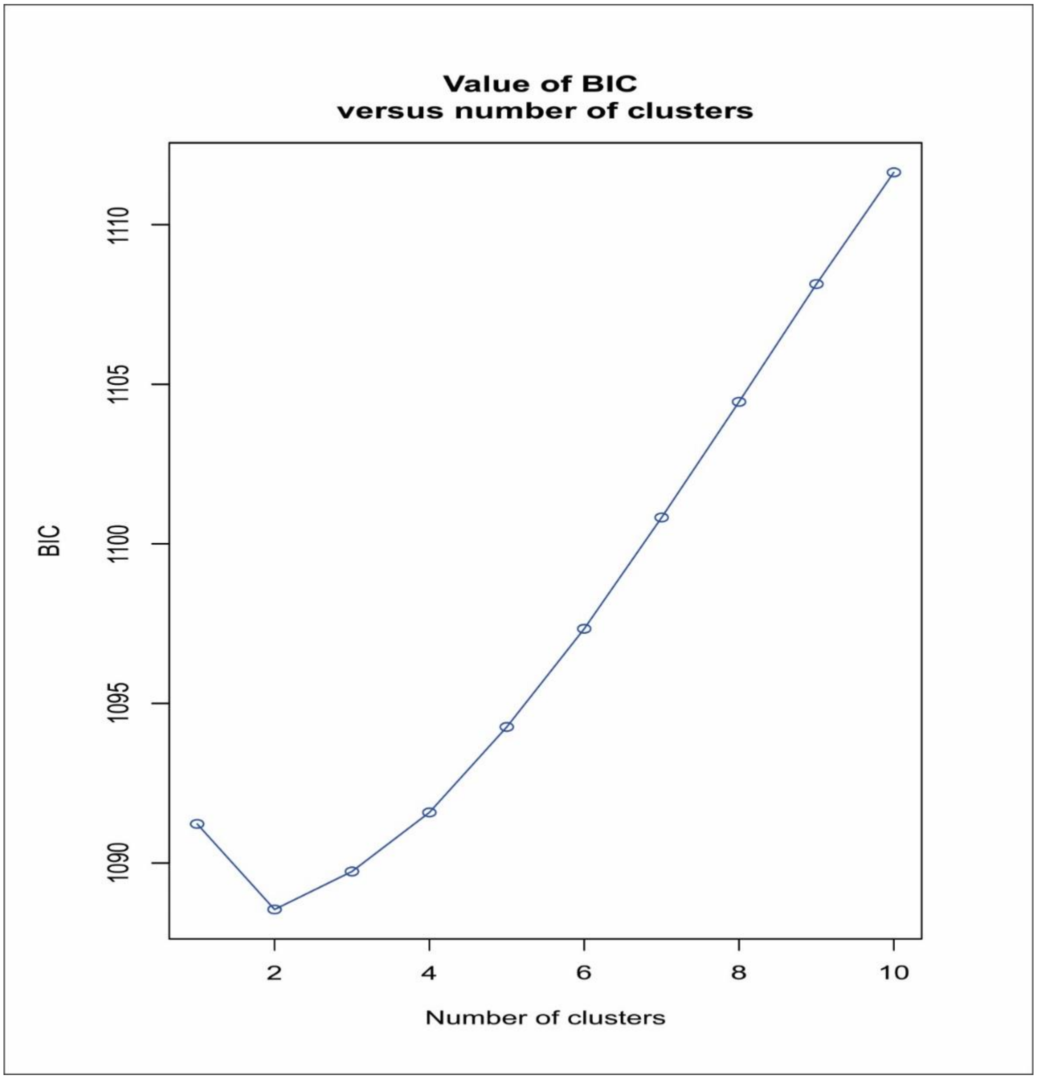


**Figure S3**. Bayesian Information Criterion (BIC) suggesting the optimal clustering solution (K=2) for the black goby dataset.
